# Association of extent of cannabis use and psychotic like intoxication experiences in a multi-national sample of First Episode Psychosis patients and controls

**DOI:** 10.1101/739425

**Authors:** Musa Sami, Diego Quattrone, Laura Ferraro, Giada Tripoli, Erika La Cascia, Charlotte Gayer-Anderson, Jean-Paul Selten, Celso Arango, Miguel Bernardo, Ilaria Tarricone, Andrea Tortelli, Giusy Gatto, Simona del Peschio, Cristina Marta Del-Ben, Bart P. Rutten, Peter B. Jones, Jim van Os, Lieuwe de Haan, Craig Morgan, Cathryn Lewis, Sagnik Bhattacharyya, Tom P Freeman, Michael Lynskey, Robin M. Murray, Marta Di Forti

## Abstract

**Aims:** First Episode Psychosis (FEP) patients who use cannabis experience more frequent psychotic and euphoric intoxication experiences compared to controls. It is not clear whether this is consequent to patients being more vulnerable to the effects of cannabis use or to their heavier pattern of use. We aimed to determine whether extent of use predicted psychotic-like and euphoric intoxication experiences in FEP patients and controls and whether this differs between groups.

**Methods:** We analysed data on lifetime cannabis using patients (n=655) and controls (n=654) across 15 sites from six countries in the EU-GEI study (2010-2015). We used multiple regression to model predictors of cannabis-induced experiences and to determine if there was an interaction between caseness and extent of use.

**Results:** Caseness, frequency of cannabis use and money spent on cannabis predicted psychotic-like and euphoric experiences, independent of other experiences (p≤0.001). For psychotic-like experiences there was a significant interaction for caseness x frequency of use (p<0.001) and caseness x money spent on cannabis (p=0.001) such that FEP patients had increased experiences at increased levels of use compared to controls. There was no similar significant interaction for euphoric experiences (p>0.5).

**Conclusions and Relevance:** FEP patients are particularly sensitive to increased psychotic-like, but not euphoric experiences, at higher frequency and amount of cannabis use compared to controls. This suggests a specific psychotomimetic response in patients related particularly to heavy cannabis use.

## Introduction

There is consistent evidence supporting an association between cannabis use and later psychosis(1). Further, patterns of cannabis use in first episode psychosis (FEP) patients are greater in terms of quantity, frequency and potency of cannabis used compared to controls from the same population(2,3). There is converging evidence that cannabis is a component cause of psychotic disorder with well-replicated evidence of dose-response effects on psychotic outcomes(3–6).

When discussing psychosis and cannabis use, it is important to differentiate between psychotic-like experiences (PEs) and clinical psychotic disorder. Clinical psychotic disorder is relatively rare whereas PEs are common and self-limiting but can be a harbinger of more serious disorder(7,8). However, the usual instruments for measuring PEs, such as the Peter’s Delusions Inventory (PDI) or the Community Assessment of Psychic Experience (CAPE), either do not specifically index drug-induced experiences as part of the intoxication state(9) or specifically exclude them(10,11).

Recreational drugs such as cannabis are used primarily for their immediate psychoactive effects. Factor analytic approaches have clustered cannabis intoxication experiences into psychotic-like experiences (cPLEs) and euphoric experiences (cEEs)(12,13). cPLEs (sometimes called psychotomimetic experiences) are worthy of study in their own right as a model for psychotic disorder. cPLEs are increased in patients versus controls(14,15); increased in those with schizotypy and those at risk of schizophrenia(13,16,17). cPLEs may predict cessation of use in a non-clinical sample(18) whereas patients with psychotic disorders report using cannabis for affect regulation and socialization, despite awareness that cannabis has a detrimental effect on positive symptoms of psychosis(19).

One study to date has reported that patients experience both cPLEs and cEEs more frequently than controls but this did not take into account increased use in patients(15). Given that both increased rates of cannabis use and increased cannabis experiences are seen in FEP, it is not yet clear how these relate to each other and whether this differs from that of controls. No study to date has examined specifically the relationship between extent of use, cannabis experiences and psychotic disorder.

We therefore studied cannabis experiences in a large international sample of FEP patients and control lifetime cannabis users. We hypothesised that: (a) we would replicate the finding of increased cPLEs and cEEs in FEP patients versus controls; (b) extent of use (as indexed by frequency of use, money spent on cannabis, and potency) would be associated with more frequent cannabis-induced experiences when adjusted for confounders; and (c) this effect would differ between cases and controls: specifically that both cPLEs and cEEs would be more affected by heavy use in FEP patients versus controls. We included THC potency as a proxy of the dose of Δ□-tetrahydrocannabinol the primary psychomimetic constituent in cannabis(20).

## Methods

The European network of national networks studying gene environment interactions in schizophrenia (EU-GEI) study is a multi-centre study comprising several work packages(21). Workpackage 2 comprises a 17 centre study across six countries (United Kingdom, Holland, Spain, France, Italy, Brazil) on first episode psychosis. Local Research Ethics Committee approval was obtained from each area.

### Sample selection

Patients and controls were recruited between May 2010 and May 2015. Patients were identified by trained EUGEI researchers across the 17 sites and invited by clinical teams to participate. For patients inclusion criteria were: (i) age 18-64; (ii) presentation with First Episode psychosis (ICD-10 F20-33); and (iii) residence within each defined locality. Exclusion Criteria were: (i) organic psychosis (ICD-10: F09); (ii) psychosis due to acute intoxication (ICD-10: F1X.5) and (iii) previous contact with mental health services for psychosis.

Controls were recruited using a quota strategy derived from local demographic data to be representative for age, sex and ethnicity of the population at risk for each site. In order to sample controls in the first instance we undertook random sampling a) from lists of all postal addresses and b) from GP lists from randomly selected surgeries. The EUGEI study aimed to over-sample certain groups (e.g. young men) using direct approaches such as local avertismenets and leaflets at local shops and community centers. Controls were excluded if they had received a diagnosis or treatment for psychotic disorder.

Further details of the EUGEI study have previously been described(22). For the purpose of this study, analysing cannabis experiences, we only analysed data from participants (both patients and controls) who reported having ever used cannabis (lifetime use).

We did not use data from two centres: Maison-Blanche (France) as this centre did not collect controls and Verona (Italy) as cannabis use data were not complete. We excluded 12 cases (1.8%) who were classified as having non-psychotic illness from the *Diagnosis and Statistical Manual IV* (DSM-IV) Operational Criteria Checklist (OPCRIT) screening of medical records.

### Measures

#### Demographics

data were collected on age, sex, ethnicity, site, country and years of education.

#### Cannabis use

A modified version of the Cannabis Experiences Questionnaire was used to collect cannabis use variables and cannabis experiences data(23). This is a researcher administrated measure which collects self-reported data on: age of first use, frequency of use (categories: every day; more than once a week; a few times a month; a few times each year; only once or twice), average money spent in a week (categories: less than €2.50; €2.50-€5.00; €5.00-€10.00, €11.00-€15.00; €16.00-€20.00; and 6 above €20). Since there is geographical variation in type of cannabis used strain data was dichotomised into ‘high potency’ preparations (THC>10%) and ‘low potency’. using published data on the expected concentration of Delta-9-tetrahydrocannabinol (THC) in the different types of cannabis available across the sites(24).

#### Other drug use

We collected data on number of other drugs used, number of cigarettes smoked per day and units of alcohol consumed daily.

#### Cannabis Experiences

Frequency of nine intoxication experiences - six cPLEs (feeling fearful; feeling crazy or mad; feeling nervy; feeling suspicious; hearing voices; seeing visions); and three cEEs (feeling happy; understanding the world better; being full of plans or ideas) were rated on a 5 point Likert scale: (0 rarely or never, 1 from time to time, 2 sometimes 3 more often than not, 4 almost always). These experiences were chosen as previous factor analytic approaches in development of the Cannabis Experiences Questionnaire showed that these experiences load significantly onto respective subscales to index psychotic-like experiences and pleasurable effects(16,25).

#### Statistical Analysis

Scores were obtained for cPLEs and cEEs by simple summation, as previously undertaken(18,23). As there were half as many euphoric experiences items as psychotic like experiences items, the scores for euphoric experiences were doubled rendering a scale of between 0 and 24 for both cPLEs and cEEs. Since such experiences can be conceptualised to index an underlying continuum both cPLEs and cEES were treated as continuous variables.

Extent of use was indexed primarily by frequency of cannabis use and by potency. In further sensitivity analysis we replaced these with money spent on cannabis use and calculated a fourth variable “frequency-potency” where we stratified frequency by use of high or low potency (i.e. categories: every day high potency; every day low potency; more than once a week high potency; more than once a week low potency; a few times a month high potency; a few times a month low potency; a few times each year high potency; a few times each year low potency; only once or twice high potency; only once or twice low potency). We calculated Pearson’s Correlation coefficients to test whether the four extent of use variables were correlated.

#### Demographics and substance use

We ascertained differences between demographic (age at assessment, sex, ethnicity, years in education, site) and cannabis use parameters (age of first use, frequency of use, money spent per week, potency, duration of use, lifetime and 12 month dependence) and other drug use parameters (cigarettes per day, units of alcohol in a day, and other drugs ever used (excluding cannabis, alcohol, tobacco and caffeine)) using t-tests for continuous variables and chi-squared for categorical variables.

### Main Analysis

We undertook to test the three hypotheses in a regression analyses framework. To test hypothesis (a) that caseness predicts experience: we regressed cannabis experiences (cPLEs and cEEs) as the dependent variables and caseness as the independent variables. To test hypothesis (b) that extent of use predicts experiences: we regressed cannabis experiences as the dependent variables and the extent of use variables as the independent variables. As the extent of use variables we entered frequency of cannabis use, and THC potency into separate models. These two variables (frequency of use and potency) were chosen to primarily index extent of use as they are both related to the extent of cannabis exposure but are distinct behaviours (for example one can use very frequently but at low potency). To test hypothesis (c): that there is an interaction between caseness and extent of use on cannabis experiences: we regressed cannabis experiences as the dependent variables and caseness and the extent of use variables alongside the interaction of caseness x extent of use. In all models we entered cPLEs as a regressor when the dependent variable was cEEs and cEEs as a regressor when the dependent variable was cPLEs to ensure that the predictors identified for relationships were independent of the other experience.

In sensitivity analyses for hypothesis (b) and (c) we ran the same regressions models using money spent on cannabis use and frequency-potency as the extent of cannabis use variables rather than the frequency or potency variables.

We undertook a further sensitivity analysis to adjust for confounders. Psychotic like experiences may be explained by a number of putative other confounders other than caseness or extent of use. We hence adjusted for firstly demographic variables (age, sex, ethnicity) in secondary models and further to this substance misuse confounders in tertiary models (number of other drugs used, tobacco use and alcohol use) as other substance misuse may arguably be related to cannabis induced experiences to see if interaction effects survived putative confounders.

cPLEs and cEEs demonstrated positive skew (cEEs 0.612, cPLEs 2.231). Because of violations of homoscedasticity in regression models we undertook all analyses using the robust regression option in STATA. For the purpose of estimation of 95% Confidence Intervals in figures (see Figure 1 & 2) we applied bootstrapping to inferential tests using 1000 samples and bias corrected and accelerated confidence intervals.

**Figure 1:**
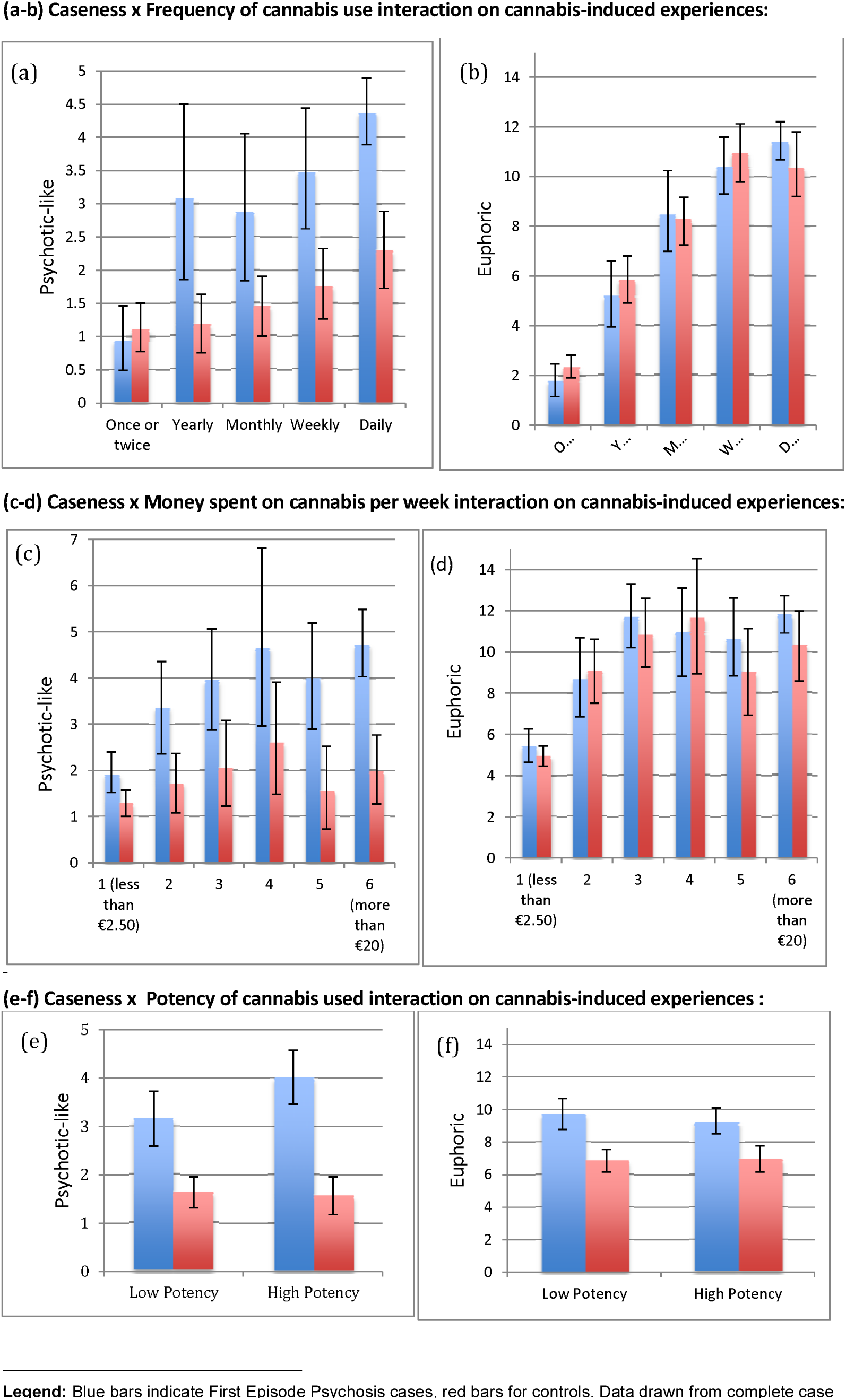
Mean cannabis-induced Psychotic-like Experiences and Euphoric Experiences scores by case and control^1^.

**Figure 2:**
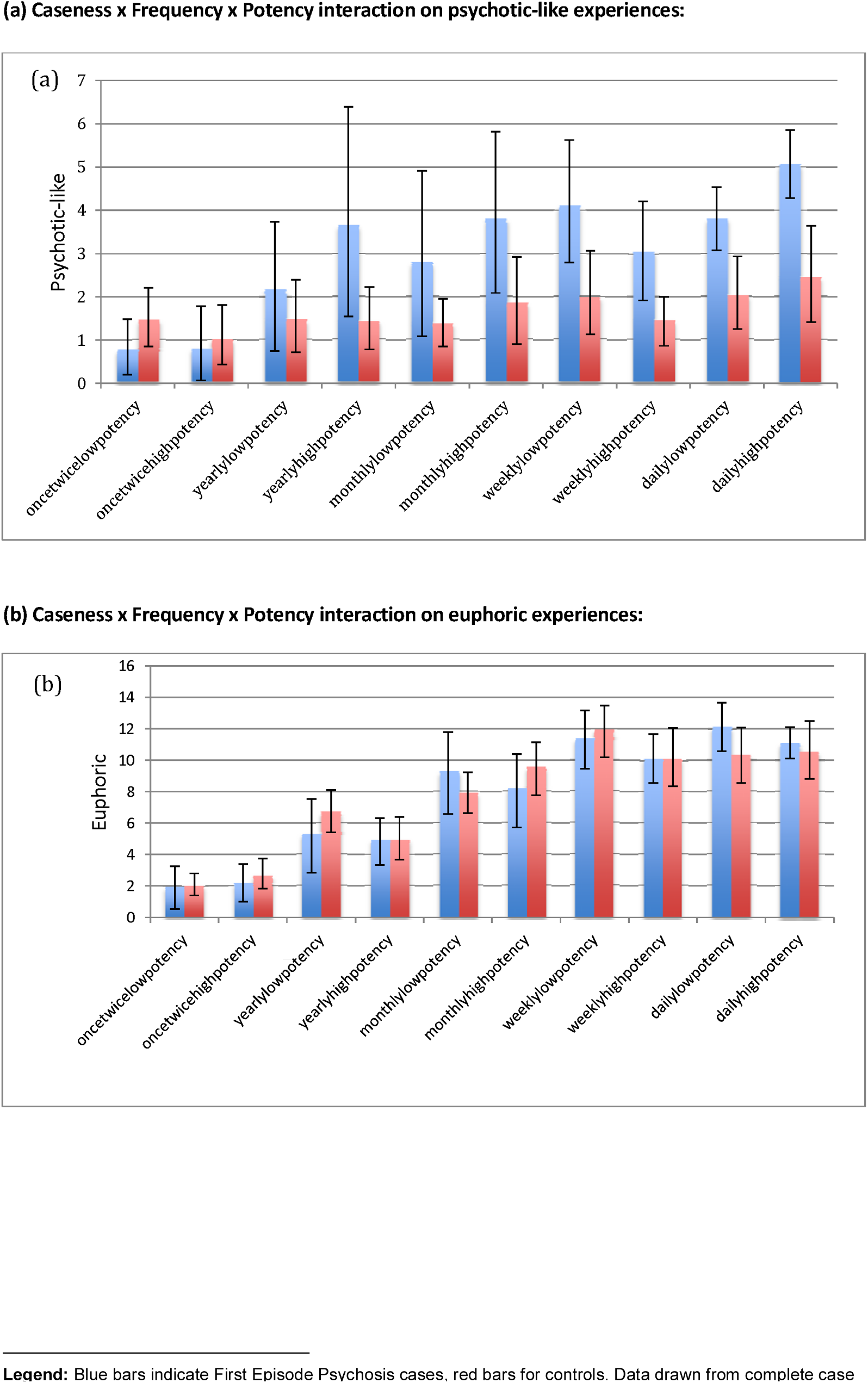
Mean cannabis-induced Psychotic-like Experiences and Euphoric Experiences scores by case and control^1^.

#### Missing data

Missing data rates are shown in Supplementary Table 4. cPLEs were available for 598/655 (91.3%) cases and 615/654 (94.0%) controls whereas cEEs scores were available for 602/655 (91.9%) cases and 616/654 (94.2%) controls.To ensure that results were not the result of systematic missing data, missing data was imputed using imputation analysis with chained equations(26) for cPLEs and and cEEs as outcome variables, independent and auxillary variables. 29 variables were included in the imputation model, including cannabis use variables (age of first use, social use, frequency, money spent, diagnosis of misuse), other drug use variables (tobacco use, alcohol use, number of other drugs used), and demographic variables (sex, age, ethnicity, site, psychosis diagnosis). Fifty datasets were imputed with 10 cycles.

Regression and main analyses were run using the imputed dataset to account for missing data. Exploratory pairwise correlation between the extent of use variables was undertaken listwise since pairwise correlation is not available using the *mi estimate* command in STATA. Data was analysed using STATA version 15.

## Results

Data were available for 1035 cases patients and 1382 controls. 655 cases (63.3% of all cases) and 654 controls (47.3% of all controls) reported ever use of cannabis and data analysis was restricted to them.

### Baseline demographics

Cases were significantly more likely than controls to be male, younger and have had fewer years of education (see Table 1a). As expected, cases were more likely to have started using cannabis younger, more likely to have used more frequently, to have used more other drugs, and smoked more cigarettes per day (see Table 1b). Detailed diagnostic, ethnicity and site data are presented in Supplementary Tables 1-3.

**Table 1a:**
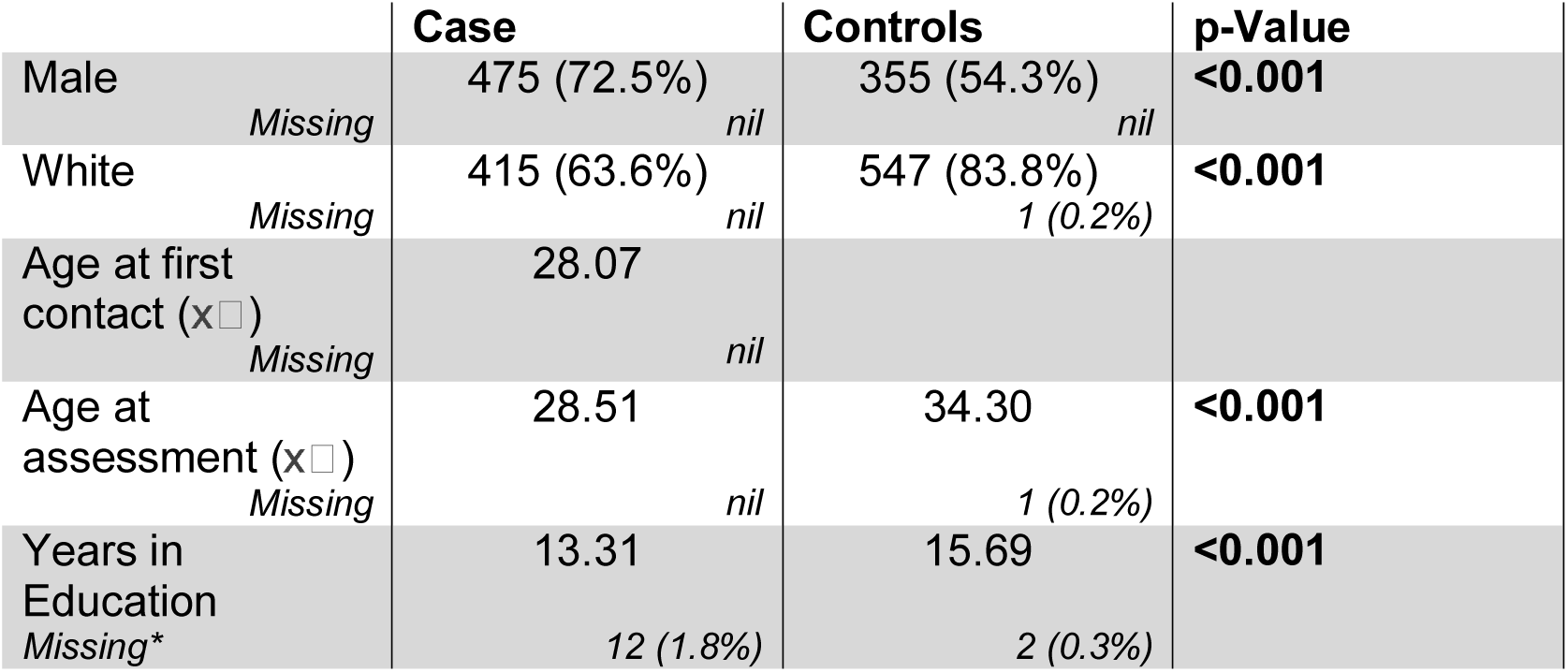
Baseline characteristics between cases and controls.

**Table 1b:**
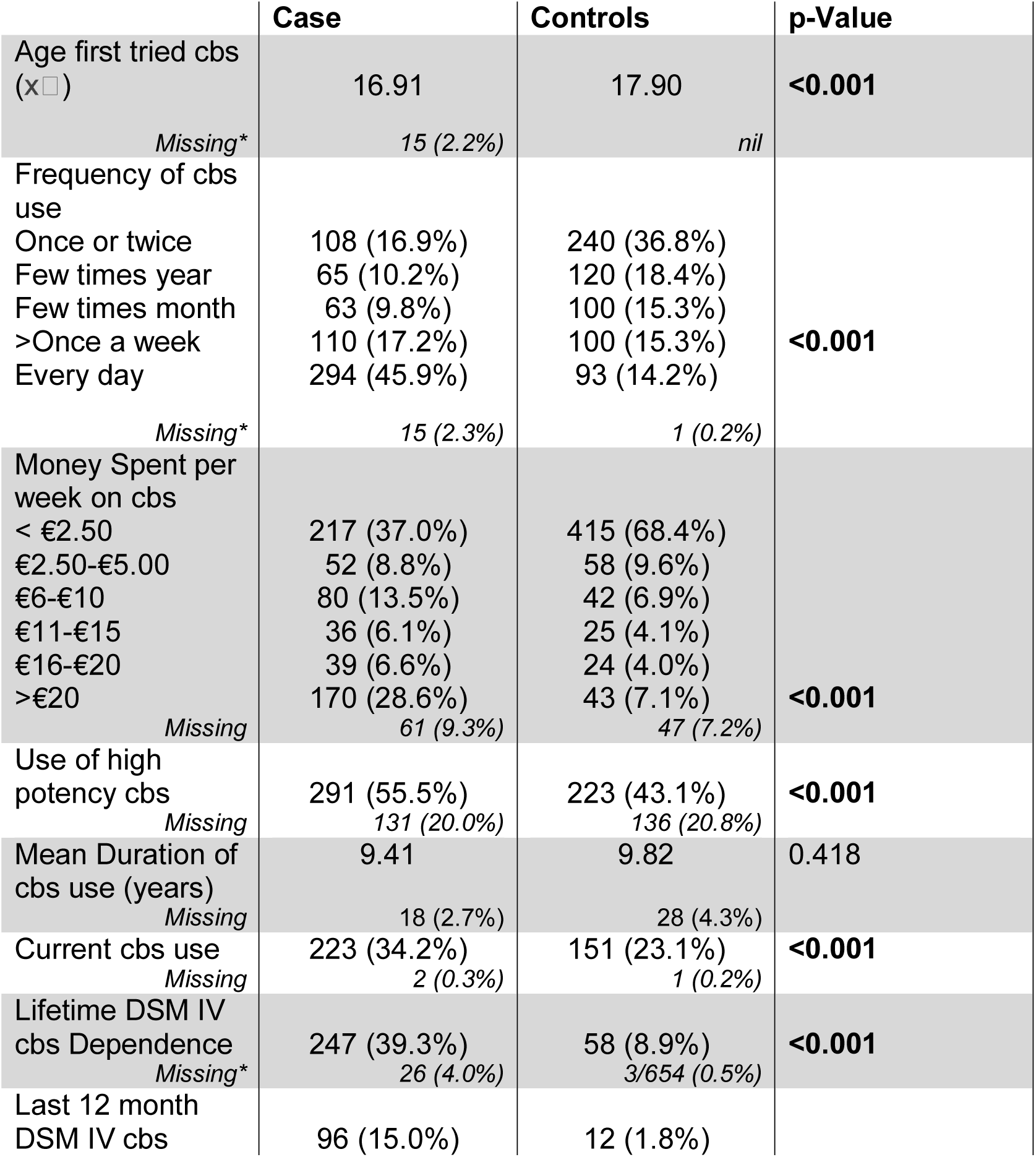
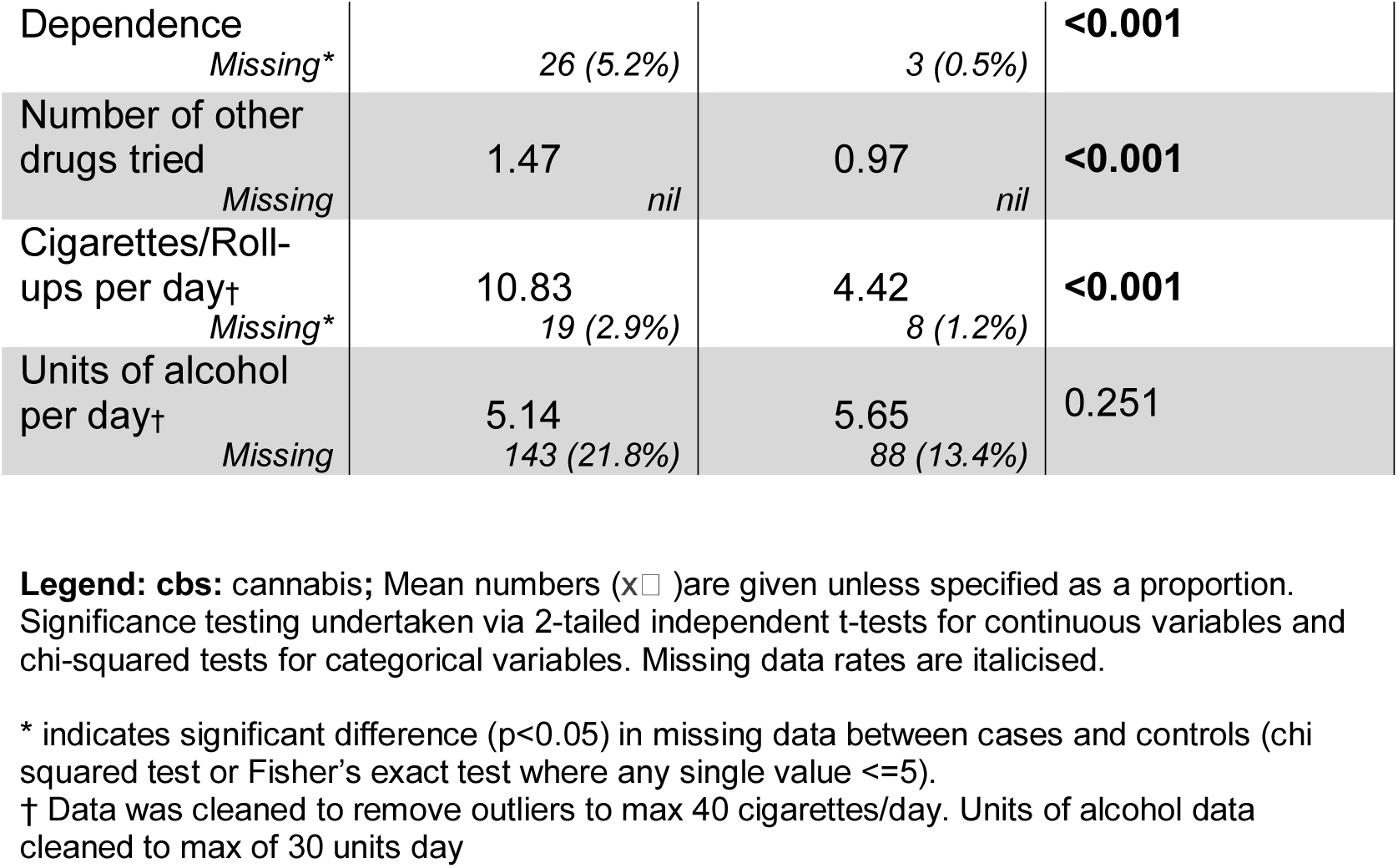
Comparison of Cannabis use patterns between cases and controls.

### Extent of use

As expected the variables indexing extent of use were significantly correlated. This was true for both the primary extent of use variables (frequency, potency) and secondary extent of use variables (money spent, frequency-potency). Frequency of use weakly correlated with dichotomised potency (r=0.121, p=0.001). Frequency of use strongly correlated with with money spent on cannabis per week (r=0.703, p<0.001) and frequency-potency (r=0.888, p<0.001) whereas potency moderately correlated with money spent on cannabis (r=0.211, p<0.001) and frequency-potency (r=0.38, p<0.001). Potency-frequency strongly correlated with money spent (p=0.727, p<0.001).

### Caseness by frequency of use on cPLEs and cEEs (hypothesis a)

As hypothesised caseness predicted cPLEs independent of cEEs (b=0.826, t=7.86, p<0.001) and predicted cEEs independent of cPLEs (b=0.840, t=4.40,p<0.001) such that patients had both more frequent psychotic-like and euphoric experiences than controls.

### Extent of use as a predictor of cPLEs and cEEs (hypothesis b)

As hypothesised extent of use predicted cPLEs independent of cEEs whether the extent of use variable was frequency of use (b=0.502, t=6.18, p<0.001), or potency (b=0.543, t=2.36, p=0.019) such that increased extent of use predicted increased psychotic-like experiences.

Similarly frequency of use predicted cEEs independent of cPLEs (b=2.17, t=21.46, p<0.001) but this was not the case with potency (b=0.210, t=0.55, p=0.58).

### Interaction Effects (hypothesis c)

Model parameters for caseness by extent of use and their interaction on predicting cannabis psychotic-like experiences can be seen in Table 2 and caseness x extent of use scores for mean experiences are shown in Figure 1 and 2.

**Table 2.** Primary Models for cannabis-induced Psychotic-Like Experiences caseness x extent of use interaction.

|  |  |  |  |
| --- | --- | --- | --- |
| **(i) Model 1 – Frequency of cannabis use as a predictor** $F(4, 1239.3)=33.65, p<0.001$ | | | |
|  | **b** | **t** | **p** |
| Frequency of cannabis use* | 0.794 | 4.74 | 0.001 |
| Caseness† | 1.354 | 6.20 | <0.001 |
| Caseness x Frequency of use‡ | 0.229 | 3.49 | <0.001 |
| Cannabis-induced Euphoric Experiences | 0.719 | 3.35 | <0.001 |
| **(ii) Model 2 – Potency of cannabis as a predictor** $F(4, 1141.9)=27.02, p<0.001$ | | | |
|  | **b** | **t** | **p** |
| Potency of cannabis* | 1.241 | 2.28 | 0.023 |
| Caseness | 0.142 | 0.42 | 0.676 |
| Caseness x Potency | 0.438 | 2.04 | 0.042 |
| Cannabis-induced Euphoric Experiences | 0.114 | 6.43 | 0.016 |
| **(iii) Model 3 – Money spent on cannabis as a predictor** $F(4, 1235.8)=33.35, p<0.001$ | | | |
|  | **b** | **t** | **p** |
| Money spent on cannabis* | 0.591 | 4.56 | <0.001 |
| Caseness | 0.267 | 1.59 | 0.112 |
| Caseness x Money spent on cannabis‡ | 0.177 | 3.29 | 0.001 |
| Cannabis-induced Euphoric Experiences | 0.084 | 4.35 | <0.001 |
| **(iv) Model 4 – Potency-frequency as a predictor** $F(4, 1241.1)=33.04, p<0.001$ | | | |
|  | **b** | **t** | **p** |
| Potency-frequency* | 0.581 | 4.09 | <0.001 |
| Caseness | 0.030 | 0.12 | 0.906 |
| Caseness x Potency-frequency‡ | 0.162 | 2.78 | 0.006 |
| Cannabis-induced Euphoric Experiences | 0.114 | 6.43 | <0.001 |

Directions of effect as follows: \*Increased extent predicts increased cPLEs; †First Episode Psychosis predicts increased cPLEs; ‡Significant caseness x extent interaction

### Caseness x frequency of use on cPLEs

There was a significant caseness effect (b=1.354, t=6.20, p=0.001); a significant effect for increased frequency of cannabis use (b=0.794, t=4.74, p<0.001); and a significant interaction between group and frequency such that increasing frequency was associated with increased difference in cPLEs between cases and controls (b=0.229, t=3.49, p=0.001).

### Caseness x potency on cPLEs

There was no significant effect of caseness (p=0.676); but an effect for potency such that increased potency was associated with increased cPLEs (b=1.241, t=2.28, p=0.023); and a significant interaction for caseness by potency (b=0.438, t=2.04, p=0.042).

### Caseness x extent of use variables on cEEs

There was evidence for increased euphoric experiences as cannabis use increased frequency (b=2.152, t=9.44, p<0.001) but not for potency (p=0.935). There was no significant interaction for either frequency or potency of cannabis use x caseness for cEEs as the dependent variable.

### Sensitivity analysis (hypothesis b)

For cPLEs results were the same when extent of use was indexed by money spent on cannabis per week (b=0.397, t=6.17, p<0.001) and potency by frequency (b=−0.412, t=6.15, p<0.001) such that both variables predicted increased psychotic-like experiences. Similarly for cEEs increased money spent on cannabis predicted cEEs independent of cPLEs (b=1.24, t=13.64, p<0.001), as did potency-frequency (b=1.38, t=14.38, p<0.001).

### Sensitivity analysis (hypothesis c)

#### Caseness x money spent on cPLEs

There was no significant effect of caseness (p=0.112); but there was a significant effect for money spent such that cPLEs increased with more money spent (b=0.591, t=4.56, p=0.001); and a significant interaction between caseness and money spent such that more money spent was associated with increased difference in cPLEs between cases and controls (b=0.177, t=3.29, p=0.001).

#### Caseness x frequency-potency on cPLEs

There was no significant effect of caseness (p=0.906); but there was a significant effect for frequency-potency such that cPLEs increased with increasing use (b=0.581, t=4.09, p<0.001); and an interaction between frequency-potency and caseness (b=0.162, t=2.78, p=0.006) such that increasing frequency-potency was associated with increased difference in cPLEs between cases and controls.

#### Caseness x extent of use variables on cEEs

There was evidence for increased euphoric experiences as cannabis use increased for money spent (b=1.109, t=5.75, p<0.001) and frequency-potency (b=1.302, t=6.33, p<0.001). There was no significant interaction for any of the extent of use variables x caseness for cEEs as the dependent variable.

### Sensitivity analysis: Adjustment for demographic and substance use covariates

In secondary models we adjusted models for cPLEs as the dependent variables for demographic covariates: the interaction terms remained significant for caseness x frequency of use (b=0.207, t=3.19, p=0.001); caseness x money spent on cannabis (b=0.163, t=3.07, p=0.002); caseness x potency (b=0.446, t=2.08, p=0.038); and caseness x potency-frequency (b=0.43, t=2.48, p=0.013). In tertiary models we additionally adjusted for substance misuse covariates: the interaction terms remained significant for caseness x frequency of use (b=0.208, t=3.23, p=0.001) and caseness x money spent on cannabus (b=0.176, t=3.30, p=0.001); caseness x potency (b=0.441, t=2.08, p=0.038); and caseness x potency-frequency (b=0.145, t=2.52, p=0.012). We conclude that the caseness x extent of use interaction for increased cPLEs for patients versus controls is robust to a number of demographic and substance use confounders.

## Discussion

To our knowledge, this represents the largest case-control study with extensive cannabis data in First Episode Psychosis ever undertaken. We (a) replicate the finding that cannabis intoxication experiences are more frequent in patients compared to controls; (b) show that extent of use as indexed by frequency of use and money spent on cannabis per week predict these experiences and (c) show that there is an interaction between caseness x frequency and caseness x money spent and caseness x frequency-potency such that increasing levels of use are associated with more frequent psychotic-like experiences (but not euphoric experiences) in patients compared with controls.

Importantly, these findings indicate that cannabis related experiences change as a function of extent of use. The Cannabis Experiences Questionnaire provides a measure of experiences as a proportion of total cannabis use, rather than a simple count of total experiences. A maximal score for cPLEs indicates that all six psychotic like experiences were experienced every time cannabis was used whereas a minimal score indicates that these experiences were never or rarely experienced, irrespective of total number of times used. Hence higher scores indicate that the experience changes rather than simply indicating an increased total number of experiences due to increased number of times that cannabis is used.

This study extends previous work(15) by showing that extent of use is a key predictor of psychotic-like experiences and that FEP patients and controls have divergent experiences with increasing extent of use. Interestingly, the same relationship does not hold for euphoric experiences as cEEs scores, when stratified by extent of use, are well-matched between cases and controls. This suggests that specific mechanisms underlie the cannabis-related increases of psychotic-like experiences which may be related to genetic predisposition and may further support a GxE interaction as has been demonstrated on cannabis use with the risk of schizophrenia spectrum disorder(27). One putative mechanism to be examined is that variation in the DRD2 and possibly AKT1 genes may render cases more likely to develop postsynaptic supersensitivity(28,29). Further work is needed to identify the specific genetic mechanisms which interact with increased extent of use.

### Strengths and Limitations

The particular strengths of this study are (i) the sample size and (ii) the international sample. The limitations include: (i) the cross-sectional design, (ii) the use of self report measures and (iii) the lack of laboratory tests of potency.

The cross-sectional design precludes interpretation about temporal sequence of associations, which means it is difficult to disentangle whether extent of use causes enhanced experience or vice-versa. Euphoric experiences (cEEs) are likely to drive use whereas this is not the case for psychotic-like experiences (cPLEs) which have previously been shown to be associated with subsequent discontinuing use(18,30). Furthermore in the case of cPLEs we included cEEs as a covariate in the model to regress out the association with euphoria. This may tentatively suggest a role for sensatisation to increasing levels of cannabis use for cPLEs in FEP.

Both exposure and outcome measures were based on self-report. There are limited methods to determine extent of use over a longer period. Hair samples can provide an estimate of use over three months, but have been shown to be unreliable in a major observational study(31). Moreover, self-report (but not hair) measures of cannabis use were found to predict acute psychotomimetic responses to cannabis(32). Additionally, self-reported data on cannabis potency is associated with its concentration of THC measured in the laboratory(33) The outcome measures, although self-reported, were based on a considerable body of work validating cannabis experiences in non-clinical, although not in clinical populations(12,23). Another limitation is that the psychotic-like experiences were rated retrospectively rather than as state measures (e.g. in an experimental design administering THC).

On the other hand, a strength of utilising retrospective self-report measures is that these are the experiences patients report to their clinicians during routine consultations. There were several differences between cases and controls, but the results persisted after adjusting for a wide variety of confounders. Perhaps most importantly cEEs were the same between patients and controls when accounted for extent of use: this indicates differences in cPLEs between FEP and controls to be specific to intrinsic biological differences between groups rather than to other confounders.

### Clinical implications

We consider this study to have a number of important findings in the clinical context. Although easily elicitable, clinicians do not routinely inquire about cPLEs in the clinical context. Our study suggests there are important differences between FEP patients and controls. Firstly our study adds to previous work(15), that patients experience cPLEs more frequently than controls. Secondly our work indicates that reduced extent of use is associated with decreased cPLEs. This is in line with evidence suggesting that FEP who continue to use cannabis, especially daily high potency experience more relapses and worse clinical outcome than those who stop after illness onset(5). Thirdly we show that FEP patients are unlikely to derive greater euphoric effects compared to controls at increased levels of use, despite more frequent psychotic-like effects. This suggests that patients and particularly those with profound cPLEs should be encouraged to reduce amount and frequency of intake; be advised that for high-potency cannabis there is limited evidence of added euphoric effect.

Taken together we have shown that extent of cannabis use is associated with enhanced psychotic-like but not euphoric experiences in First Episode Psychosis patients compared to controls. Further research should aim to determine the biological mechanism underpinning differences between patients and controls.

## Funding

This work was supported by the European Community’s Seventh Framework Programme under grant agreement No. HEALTH-F2-2010-241909 (Project EU-GEI). The Brazilian study was funded by the São Paulo Research Foundation under grant number 2012/0417-0. Dr Marta Di Forti is funded by the MRC not involved in design and conduct of the study; collection, management, analysis and interpretation of the data; preparation, review or approval of the manuscript, and decision to submit the manuscript for publication. Dr Sami is supported by a Medical Research Council Clinical Research Training Fellowship (MR/P001408/1). Dr Freeman was supported by a Senior Academic Fellowship from the Society for the Study of Addiction.

Dr Marta Di Forti and Dr Musa Sami had full access to all the data in the study and take responsibility for the integrity of the data and the accuracy of the data analyses.

