## Supplement for "Association of extent of cannabis use and psychotic like intoxication experiences in a multi-national sample of First Episode Psychosis patients and controls"

**Online supplement:**

| **sTable 1 DSM IV diagnosis of cases* (Operational Criteria Checklist (OPCRIT):** \| ***n*** \| *%*; No diagnosis \| 69 \| 10.3; Major depressive disorder with psychosis \| 56 \| 8.4; Manic episode with psychosis \| 58 \| 8.7; Schizophrenia \| 179 \| 26.8; Schizophreniform disorder \| 115 \| 17.2; Schizoaffective disorder, depressive type \| 13 \| 1.9; Schizoaffective disorder, bipolar type \| 19 \| 2.8; Delusional disorder \| 23 \| 3.4; Psychosis not otherwise specified (atypical psychosis) \| 95 \| 14.2; Bipolar I disorder \| 20 \| 3.0; *Missing* \| 8 \| 1.2; **Total** \| **655***^†^ \| **98.2***^†^ *Ever cannabis using patients with FEP ^†^ Cases excluded from analysis (not shown in table above) were those with diagnosis of non psychotic disorders (n=12; 1.8%): Moderate Major depressive disorder; Major depressive disorder severe; hypomanic episode; Manic episode without psychosis **sTable 2: Cases & Controls by site:** | | | | |
| --- | --- | --- | --- | --- |
|  | |  | | **Total** |
|  |  | **Case** | **Control** |  |
| **Brazil**  Ribeirão Preto |  | 89 (13.6%) | 60 (9.2%) | 149 (11.4%) |
| **France**  Val-de-Marne (Paris)  Puy-de-Dôme (Clermont-Ferrand) |  | 31 (4.7%) | 38 (5.8%) | 69 (5.3%) |
|  |  | 8 (1.2%) | 23 (3.5%) | 31 (2.4%) |
| **Holland**  Amsterdam  Gouda and Voorhout |  | 86 (13.1%)  69 (10.5%) | 64 (9.8%)  57 (8.7%) | 150 (11.5%)  126 (9.6%) |
| **Italy**  Bologna  Palermo |  | 35 (5.3%)  37 (5.6%) | 39 (6.0%)  59 (9.0%) | 74 (5.7%)  96 (7.3%) |
| **Spain**  Barcelona  Cuenca  Galicia  Madrid  Oviedo  Valencia |  | 22 (3.4%)  13 (2.0%) | 23 (3.5%)  19 (2.9%) | 45 (3.4%)  32 (2.4%) |
|  |  | 19 (2.9%)  29 (4.4%)  21 (3.2%)  29 (4.4%) | 24 (3.7%)  19 (2.9%)  21 (3.2%)  16 (2.4%) | 43 (3.3%)  48 (3.7%)  42 (3.2%)  45 (3.4%) |
| **United Kingdom**  Cambridge  London |  | 29 (4.4%)  138 (21.1%) | 51 (7.8%)  141 (21.6%) | 80 (6.1%)  279 (21.3%) |
| **Total** |  | **655 (100.0%)** | **654 (100.0%)** | **1321 (100.0%)** |

**sTable 3: Cases & Controls by ethnicity**

|  | | |  | | | **Total** |
| --- | --- | --- | --- | --- | --- | --- |
|  |  |  | **Case** | | **Control** |  |
|  | White |  | 415 (63.4%) | | 547 (83.8%) | 962 (73.5%) |
|  | Black |  | 105 (16.0%) | | 46 (7.0%) | 151 (11.5%) |
|  | Mixed |  | 60 (9.2%) | | 31 (4.7%) | 91 (7.0%) |
|  | Asian |  | 22 (3.4%) | | 12 (4.7%) | 34 (2.6%) |
|  | North African |  | 31 (4.7%) | | 7 (1.1%) | 38 (2.9%) |
|  | Other |  | 22 (3.4%) | | 10 (1.5%) | 32 (2.4%) |
|  | **Total** | |  | **655 (100.0%)** | **653 (99.8%)*** | **1308 (99.9%)** |

*Ethnicity data for one control was missing.

**sTable 4: Missing data rates**

|  | **Case** | **Controls** |
| --- | --- | --- |
| Male | *nil* | *nil* |
| White | *nil* | *1 (0.2%)* |
| Age at first contact | *nil* |  |
| Age at assessment | *nil* | *1 (0.2%)* |
| **Years in Education** | ***12 (1.8%)*** | ***2 (0.3%)*** |

| **Age first tried cbs** | ***15 (2.2%)*** | ***nil*** |
| --- | --- | --- |
| **Frequency of cbs use** | ***15 (2.3%)*** | ***1 (0.2%)*** |
| Money Spent per week on cbs | *61 (9.3%)* | *47 (7.2%)* |
| Use of high potency cbs | *131 (20.0%)* | *136 (20.8%)* |
| Mean Duration of cbs use (years) | 18 (2.7%) | 28 (4.3%) |
| Current cbs use | *2 (0.3%)* | *1 (0.2%)* |
| **Lifetime DSM IV cbs Dependence** | ***26 (4.0%)*** | ***3/654 (0.5%)*** |
| **Last 12 month DSM IV cbs Dependence** | **26 (5.2%)** | **3 (0.5%)** |
| Number of other drugs tried | *nil* | *nil* |
| **Cigarettes/Roll-ups per day** | ***19 (2.9%)*** | ***8 (1.2%)*** |
| Units of alcohol per day | *143 (21.8%)* | *88 (13.4%)* |
| cPLEs | 57 (8.7%) | 39 (6.0%) |
| cEEs | 53 (8.1%) | 38 (5.8%) |

**Legend: cbs:** cannabis**;** Bold typeface indicates significant difference (p<0.05) in missing data between cases and controls (chi squared test or Fisher’s exact test where any single value <=5).
